# DNA Methylation is stable after MMEJ and NHEJ double strand break repair

**DOI:** 10.1101/2022.07.21.500965

**Authors:** Ester Bnaya, Shay Shilo, Tzah Feldman, Tal Bacharach, Yoni Moskovitz, Aditee Kadam, Paz Yedidim, Hadas Azogy, Nathali Kaushansky, Noa Chapal-Ilani, Liran I. Shlush

**Author notes:** These authors contributed equally to this work (Ester Bnaya, Shay Shilo). Corresponding author - Liran I. Shlush.

## Abstract

DNA double strand breaks (DSBs) are a major source of mutations. Both non-homologous-end-joining (NHEJ) and microhomology-mediated-end-joining (MMEJ) DSB repair pathways are error prone and produce deletions, which can lead to cancer. DSBs also lead to epigenetic changes, including demethylation, which is involved in carcinogenesis. Of specific interest is the MMEJ repair pathway, as it requires methylation restoration around the break, as a result of the resection and formation of single stranded (ssDNA) intermediates. While, methylation patterns after homologous recombination (HR) have been partially studied, the methylation status after MMEJ and NHEJ remains poorly reported, and can be relevant for cancer. To study methylation patterns around DSB after NHEJ and MMEJ repair, we used targeted bisulfite-sequencing (BS-seq) to quantify methylation of dozens of single cell clones after induction of DSB by CRISPR. Each single cell clone was classified according to the sequence signature to a specific repair mechanism: NHEJ or MMEJ. Comparison of single cell clones after DSB to control cells, without DSB, demonstrated correct restoration of the methylation levels. No difference in methylation patterns was noticed when comparing NHEJ to MMEJ. Methylation levels in gene body, highly methylated CpGs (n=61, 4000 base pairs around DSB) and in low methylation CpGs (n=19), remained stable after both MMEJ and NHEJ. Gene body methylation persisted even on the background of *DNMT3A* R882C mutation, the most prevalent preleukemic mutation, in which the *de novo* methylation machinery is compromised. An exception observed in a single CpG site (*ASXL1* 995) which demonstrated elevated methylation rate after DSB repair only in the presence of WT *DNMT3A*. In summary, DNA methylation restoration demonstrated high fidelity after DSB both in methylated and unmethylated gene body, even in cases where DNA resections and deletions occurred.

**Author Summary:** DNA holds the genetic information. Modifications on the DNA molecule such as DNA methylation are crucial for the genetic regulation. DNA damage is harmful to the cell and needs to be repaired. Different repair mechanisms may result in mutations that can be identified according to their sequence signature. In the work presented here we examined changes in DNA methylation after different types of DNA repair, MMEJ and NHEJ. We found that DNA methylation is highly stable after DNA repair regardless to the repair mechanism and genomic context.

## Introduction

### DSB repair

DNA DSB repair is an essential process for cell survival. Upon DSB formation, cells have several possible repair pathways. A common classification divides the repair mechanisms according to whether or not it uses the homologous chromosome as a template. Homologous recombination (HR) repair pathways include the crossover (CO) and non-crossover (NCO), also known as gene conversion (GC) or synthesis dependent strand annealing (SDSA). Those usually result in a copy neutral loss of heterozygosity (CN-LOH) [1,2]. The other class of repair pathways is based on an intra-chromosomal repair and can be identified according to sequence signatures, mostly small to medium deletions. Those include NHEJ, MMEJ and single-strand annealing (SSA) [3]. While NHEJ use none or minimal sequence homology, MMEJ and SSA are based on larger intra-chromosomal homology (>5 and >30 bp, respectively) [3–5]. All the homology-based pathways initiate with resection of the 5’ strand to enable pairing of the sequences and initiate template base polymerization [4].

### Long-term epigenetic changes result of DSB repair

The epigenetic landscape is in an extensive interplay with the repair mechanism. The contribution of the epigenetic landscape to the repair pathway of choice [6–11] and the short-term changes to the epigenetic landscape of [12–14] are well established. However, only a handful of studies addressed the subject of long-term epigenetic changes following DSB repair.

DNA methylation pattern is inherited to the daughter cells [15] and therefore is a leading candidate to mediate long-term epigenetic change following DSB repair. The MMEJ repair is in particular susceptibility for loss of methylation, since during MMEJ-repair, the 5’ strand is removed during resection and therefore the DNA methylation is erased and needs to be restored [4]. It is assumed that the DNA methyl-transferase *DNMT1*, which can be active on hemimethylated DNA might be responsible for methylation repair after MMEJ [15–17]. However, methylation changes as result of DSB or base excision repair were also found to be mediated by other DNA methyl-transferases, such as *DNMT3A* and *DNMT3B* [15,16,18–20].

Most previous studies focused on the interaction between HR and methylation. In plants, no methylation changes were noticed after NHEJ [21], but methylation loss was observed after HR during gene targeting [22]. In mammals, comparison of construct added into two mice genomes through HR and NHEJ revealed increase of methylation associated HR but not with NHEJ [23]. In human cells, it has been demonstrated that the flanking regions of a DSB repaired by HR can undergo *de novo* methylation by *DNMT1* with involvement of *DNMT3A* resulting in gene silencing [19,20]. The methylation changes after HR were shown to be transcription-dependent and occur in 15-20 days after HR [20].

As the cumulative sum of DSB increase with age [24] and they must be repaired, it is speculated that aging stem cells hold in their genomes tens of DSB repaired sites. It remains unclear and controversial whether DSB repair induces gain of methylation [25], mediates a mechanism for passive DNA demethylation [15] or is it dependent on the repair mechanism [23]. If so, it raise the question: do those epigenetic scars of these breaks remain in the aging stem cell genome?

### MMEJ and methylation in myeloid malignancies

Myeloid malignancies are a relevant system to learn the effect of methylation changes as result of DSB since myeloid malignancies and the preleukemic state of clonal hematopoiesis (CH) are enriched with mutations in epigenetic regulators [26–28]. Recently, we demonstrated that the most common deletions in myeloid malignancies and CH are the result of MMEJ repair. A CRISPR based system induced the formation of the naturally occurring recurrent deletions in *ASXL1, SRSF2* and *CALR* together with small indels, which are the result of NHEJ repair [29]. To study methylation patterns accurately after different DSB repair pathways we created single cell clones following DSB, so that all the cells in the clone underwent the same repair. Due to large variability in the methylation background of single cells [30,31], large number of single cell clones were generated, both after DSB and the controls without DSB. To examine the epigenetic implications of MMEJ and NHEJ, methylation was quantified in the flanking regions of the DSB in a set of single cell clones and no DSB single cell control cells.

## Results

### Epigenetic stability after DSB in highly methylated gene body (*ASXL1)*

First, we aimed at characterizing the epigenetic changes after DSB in a gene body known to carry homozygous methylated regions. For this aim, we chose to study the methylation level at the flanking regions (∼2 kbp up and downstream) of the most common site of MMEJ and NHEJ in acute myeloid leukemia (AML), *ASXL1* exon 12. We first studied bulk methylation in K562 cells that were not edited in the *ASXL1* region (controls) (Fig. 1A). The region exhibited mostly high levels of methylation, consistent with homozygote gene body methylation (S1 Fig).

**Fig 1.**
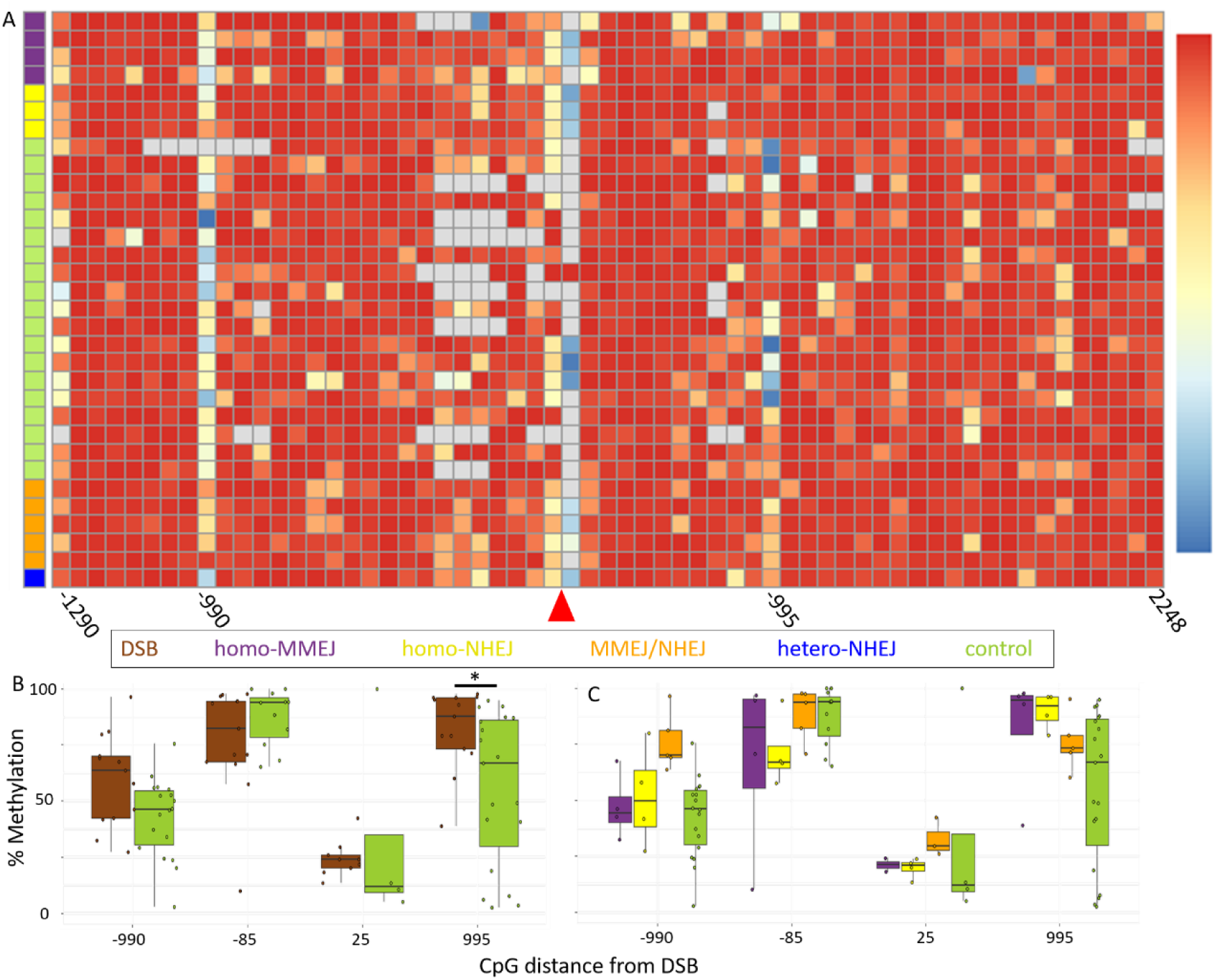
Epigenetic scars after DSB in highly methylated gene body (*ASXL1*). Measurement of DNA methylation change after DSB in the *ASXL1* gene, assessed by DNA bisulfite sequencing of homo-MMEJ (purple), homo-NHEJ (yellow), mono-NHEJ (blue), MMEJ/NHEJ (orange), and control (green) clones. **(A)** Methylation levels were scaled from 0 (absence of methylation - blue) to 100% (complete methylation - red, missing data - gray) (methods). CpG coordinates (X-axis, S4 table) are in relation to the DSB site (red triangle). Samples classification are color coded according to the mechanism of repair on the left of each row. Comparison of methylation levels in four most variable CpG sites of **(B)** all DSB outcomes (brown) or **(C)** according to a specific repair mechanism. MMEJ (purple), NHEJ (yellow), MMEJ/NHEJ (orange), and controls (green). The asterisk indicate a significant difference (<0.05) between the DSB and the control groups in position 995. Methylation is the ratio between the C and T after BS conversion (methods). All comparisons were performed using a two-tailed, non-paired, nonparametric Wilcoxon rank sum test with 95% confidence interval and FDR multiple hypothesis correction.

To induce DSB in a controlled manner, we used the CRISPR/CAS9 system that recapitulates the naturally occurring deletions in *ASXL1* [29]. After introducing DSBs, single-cell clones were established. Two types of single cell clones were used as control: non-edited K562 single cell clones, and single cell clones, which were exposed to CRISPR/CAS9 but with a target on a different chromosome (S1 Table). The methylation level at the single cell clone level was measured by targeted bisulfite sequencing (BS-seq) next generation sequencing (NGS) to 61 CpG site.

The methylation in the single cell controls found to be consistent between the two single cell clone control groups (Fig. 1A). An intriguing insight emerged when the single cell clones were compared to the bulk control. Few CpG sites which exhibited lower levels of methylation in the bulk demonstrated a variable and dynamic trend of methylation at the single cell clone level, not a constant reduced level of methylation (S1 Fig).

Targeted DNA NGS around the site of the DSB, was used to define the different types of repair in each single cell clone (S2-3 Table). MMEJ repair was defined by a deletion of a sequence flanked by homologous sequences including a deletion of one of the homologies. NHEJ was defined as small insertions or deletions (indels) of less than 5 bp at the DSB site. The repair occurred in a homozygous or heterozygous manner according to the triploid nature of the K562 cell line [32]. The single cell clones were classified to the following groups: homozygote MMEJ, homozygote NHEJ (small indel, not necessarily the same indel on the different alleles), MMEJ or NHEJ in one allele and no detectable change on the others (heteroMMEJ or heteroNHEJ), and heterozygosity of NHEJ and MMEJ (MMEJ/NHEJ) (S3 Table, Fig. 1A, C). The different outcomes of the DSB repair were consistent with previous results from the same system [29].

The methylation state associated with the repair mechanism of each clone was measured in the 61 CpG sites up and downstream to the DSB (∼2kb from each side) (S4 Table). Despite methylation variability in several sites, most of the methylation sites were stable when comparing single cell clones after DSB repair to controls (Fig. 1B), or when comparing single cell clones after MMEJ to controls or to clones after NHEJ (Fig. 1C). One exception was the CpG site 995 which demonstrated a significant (p<0.05) increase in methylation in comparison to the highly variable basic state in the single cell control clones. These results suggest that cells maintain and restore the level of methylation despite the genetic changes following the DSB. Furthermore, the methylation machinery restores the loss of methylation in the newly synthesized strand after MMEJ repair.

### Methylation restoration after DSB is stable on *DNMT3A* R882C background

The accurate restoration of methylation after DSB we observed in most of the CpG sites in K562 cells in *ASXL1* (Fig. 1) is most probably related to the rapid recruitment of *DNMT1* to the DSB site regardless of the cells cycle state of the cells [33]. However, the increase in methylation at the most variable CpG site (995) may indicate an involvement of dynamic processes. As other DNA methyl transferases (DNMTs) were found to be associated with DNA repair machineries [19,20,25,34,35], we examined whether DSB may provoke methylation scar by mutated DNMTs. *DNMT3A* R882C mutation is a common AML mutation in a DNA methylation regulator [36].

To examine possible influence of the mutation on methylation maintenance and restoration after DSB, we used the OCI-AML3 cell line carrying the recurrent *DNMT3A* R882C mutation [37]. We induced DSBs, grew single cell clones and classified the repair of *ASXL1* in the same way as done for the K562 cells. Methylation analysis of 56 of the CpG positions in the control clones demonstrated that even on the background of *DNMT3A* R882C mutation the basic high levels of methylation were stable. The methylation level in the flanking sequences of the DSB in *ASXL1* provided evidence that DSB did not introduce any methylation changes even under *DNMT3A* R882C mutation background (Fig. 2A, S2 Fig A-B, S5 Table).

**Fig 2.**
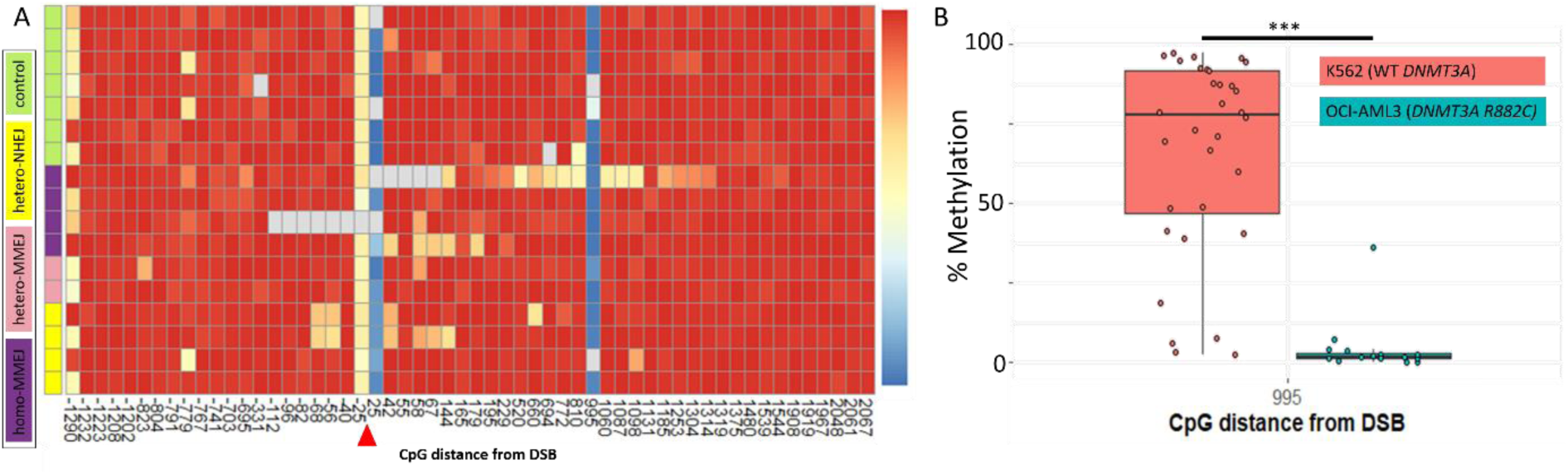
Methylation stability in OCI-AML3 single cell clones following DSB in *ASXL1*. Measurement of DNA methylation changes after DSB in *ASXL1* gene on the background of *DNMT3A* R882C, assessed by DNA bisulfite sequencing of homo-MMEJ (purple), hetero-MMEJ (pink), hetero-NHEJ (yellow), and control (green) clones. **(A)** Methylation levels were scaled from 0 (blue) to 1 (red) (missing data - gray). CpG coordinates (X-axis) are in relation to the DSB site (red triangle). DSB repair outcome for each sample is indicated at the left of each row. **(B)** Significant difference (p<0.0001) in basal methylation levels of CpG site at position 995 bp downstream to the DSB site between K562 with WT *DNMT3A* (pink) and OCI-AML3 with R882 *DNMT3A* (turquoise).

Nonetheless, while exhibiting stability in most of the methylation landscape, a deviation was observed between K562 and OCI-AML3 with R882C background. The basic level of methylation of CpG at position 995, the most variable CpG site of the control clones in K562 cell, exhibited on the background of R882C a complete loss of methylation (p<0.0001) (Fig. 2D). Moreover, apart from the K562 cells which demonstrated an elevated methylation as result of DSB repair at the CpG position 995, the induction of DSB repair did not change the level of methylation at the position and it remained completely unmethylated (Fig. 2A).

Overall, all the single-cell clones, regardless to the repair mechanism, presented restoration of the methylation patterns (Fig. 2B-C, S3, 5 Tables), except for CpG site 995. This finding is a demonstration of the robustness of the methylation restoration after DSB even on the background of an aberrant methylation machinery, the *DNMT3A* R882C mutation.

### Epigenetic stability after DSB in the unmethylated *SRSF2* gene in K562 cells

Since we noticed a robust restoration of methylation in highly methylated regions in both cell lines accompanied with elevated methylation at CpG 995 in K562, we examined how the repair machinery preform on an unmethylated region in K562 cells. Is it maintaining the correct level of methylation or whether the DSB induces gain of methylation, as observed in position 995 in K562 and was reported in the past [19,20].

To address this question, we chose to focus in the unmethylated *SRSF2* gene in K562 cells. *SRSF2* plays an important role in pathogenesis of AML and was part of the MMEJ induction system established previously [29,38]. The high density of CpG sites around the canonical MMEJ deletion in *SRSF2* limited our ability to explore the region directly with targeted BS-seq. Instead, a region containing 19 CpG sites approximately 200 bp downstream to the canonical *SRSF2* deletion was targeted (Fig. S3). The targeted region resides over coding and non-coding regions of the *SRSF2* transcript. This region tests the methylation status downstream to the DSB. To adjust for possible changes upstream to DSB, two additional gRNAs were used to induce DSB downstream to the region where the methylation is measured. In this configuration, the CpG sites will be ∼100 and ∼400 bp upstream to the DSB. Those two additional DSB sites were in the *SRSF2* promoter and one of them overlaps with the gene body of the downstream gene *MFSD11* (Fig. S3). We validated the gRNA from the recently described system [29] and the two additional gRNAs for their ability to induce MMEJ deletions, including the canonical recurrent *SRSF2* deletion, and NHEJ repair. All the examined gRNAs were found to induce both MMEJ and NHEJ (Fig. S3, S3 Table).

First the CpG targets in *SRSF2* were examined for their methylation level. The great majority of CpG sites were unmethylated (∼0% methylation) (Fig. 3A). Four sites exhibited a variable pattern of methylation. No gain of methylation was observed in the unmethylated sites after DSB repair with or without regard to the type of repair (S3, 6 Tables, Fig. 3A-C). Intriguingly, one of the four methylated sites demonstrated a slight but consistent elevation in methylation in all the DSB repair outcomes, regardless to the mechanism of repair (p<0.001). This indicates that although gain of methylation is possible in the region and according to the cellular state, the correct restoration of methylation is performed and not the addition of methylation indifferently after induction of DSB.

**Fig 3.**
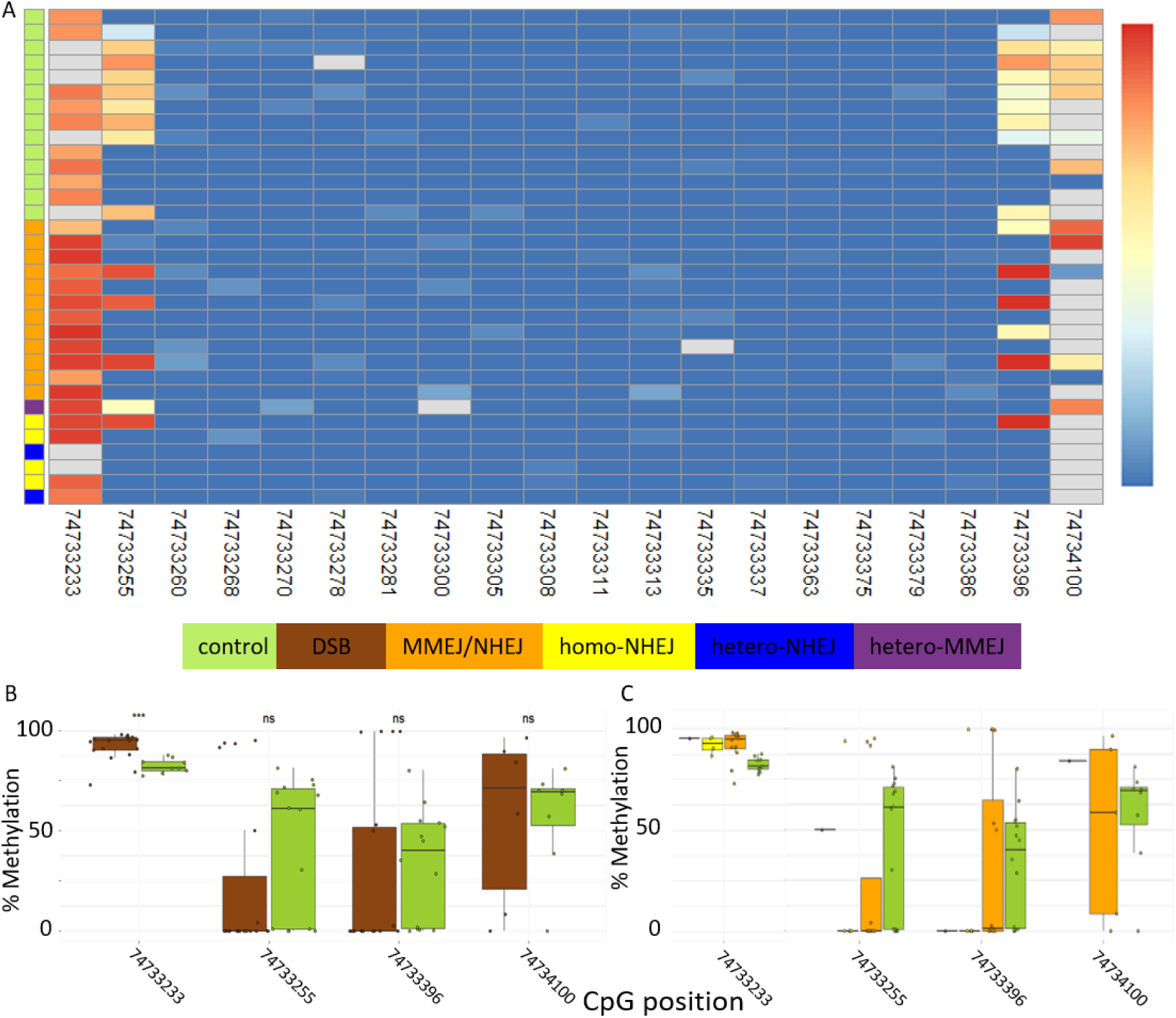
Epigenetic landscape after DSB in the unmethylated *SRSF2* gene in K562 cells. DNA methylation measurements after DSB in *SRSF2* gene, assessed by DNA bisulfite sequencing of homoMMEJ (purple), homoNHEJ (yellow), monoNHEJ (blue), MMEJ/NHEJ (orange), and control (green) clones. **(A)** Methylation levels were scaled from 0 (blue) to 1 (red) (gray - missing data). The CpG sites are ordered according to their genomic coordinates (X-axis). Comparison of methylation levels of highly variable selected CpG sites. **(B)** All DSB induced outcomes (brown) to controls (green), **(C)** according to the repair mechanism, MMEJ (purple), NHEJ (yellow), MMEJ/NHEJ (orange) and controls (green). All comparisons were performed using a two-tailed, non-paired, nonparametric Wilcoxon rank sum test with 95% confidence interval and FDR multiple hypothesis correction.

## Discussion

In the current study, we tested the cell ability to restore DNA methylation patterns that could potentially be altered after MMEJ and NHEJ DSB repair. Although MMEJ and NHEJ are error prone, our findings provide evidence that after DSB the DNA methylation is less promiscuous in comparison to the DNA sequence. While we could create many single cells with either small indels indicative of NHEJ repair and larger indels of a sequence surrounded by micro-homologies due to MMEJ repair, we could not observe any consistent methylation changes after DSB repair regardless of the repair mechanisms, or the original methylation status of the region (hypo/hyper methylated). Only a single site out of the 80 sites examined demonstrated a significant and transition from low to high methylation.

Our results prove that the overwhelming majority of DNA methylation does not change significantly after MMEJ or NHEJ repair. The field of methylation changes as a result of DSB is ambiguous with mixture of results indicative for gain, loss, or maintenance of methylation after DSB repair [22,23,25,39]. These inconsistencies might result from the different organisms, tissues, or generations after DSB induction or from difference in repair mechanisms. Such discrepancies might also be related to the complexity of the DNA repair machinery and its interaction with methylation. Thus in the current study to properly address the contribution of DSBs to methylation, the following parameters were be carefully controlled: 1) To study the effect of different repair machineries on methylation the research was done on single cell clones. Each clone represented a specific repair pathway, DNA signature, defined by sequencing. 2) To account for epigenetic polymorphisms, stochastic formation of differentially methylated regions, and dynamic changes in methylation the background methylation patterns in uncut control cells was done on tens of single cell clones. 3) The effect of DSB repair on methylation was tested in different regions of the genome with different ground state, methylated vs. unmethylated gene bodies. 4) The cells were tested at the cellular state the DSB was formed to avoid methylation changes as a result of differentiation and reprograming, as might happen in studies that are conducted in different generations. 5) The measurement of the methylation was done with quantitative method, targeted next generation BS-seq, and not Sanger BS Sanger sequencing or methylation chips that are noisy and average the signal.

Specifically, in MMEJ the restoration of the methylation after resection is accompanied with gap filling. This process is considered to be accurately mediated by Polθ, but other polymerases with variable accuracy may also be involve in the process [3,29,40]. While DNA sequence is stable, methylation of the DNA is more dynamic [30,31]. Indeed, we observed sites with variable methylation in the gene body and promoter regions even in the ground state of the control group (S1 Fig.). This indicates that methylation is not only restored but also continuously maintained. The CpG site 995 downstream to the *ASXL1* DSB site is an example for a site that is constantly maintained (Fig. 2D). On the background of wild type *DNMT3A* it is highly variable even without the induction of DSB, indicating for a dynamic process. However, under *DNMT3A* R882C mutation the site is completely unmethylated (Fig. 2A, D).

As DSBs are prominent in leukemia etiology and DNA methylation regulators are damaged in many myeloid leukemia and preleukemic CH [41], we examined whether mutations in *DNMT3A* might alter methylation reconstitution after DSB. Our results provided evidence that the reconstitution of methylation after MMEJ or NHEJ DSB repair is not influenced by the most common *DNMT3A* mutation. DSBs are known to induce histone modifications [12–14], our results suggests that such epigenetic changes do not increase changes in DNA methylation.

However, *DNMT3A* R882C mutation affect the *ASXL1* CpG site 995. This was the only incident with significant gain of methylation, and this phenomenon was abolished on the background of R882C mutation. Cells harboring the mutation, OCI-AML3 had complete loss of methylation in the site in comparison to a variable methylation on WT background or high methylation in WT background after DSB. The generalization of a conclusion based on a single site is impractical and should be examined in a genome wide context of sites affected by R882C mutation in the single cell clone level.

While in the current study we were able to separate different repair pathways and included large number of uncut single cell clones and bulk analysis, our studies are limited to only two regions of the genome, methylated and unmethylated, and only MMEJ and NHEJ. While these regions are of interest to the AML community as they are hotspots for DSBs among leukemia patients, it remains unclear whether other regions of the genome will behave in a similar way, and whether methylation is dynamic after HR. Furthermore, the current study is underpowered to identify low frequency methylation changes after DSB repair.

Altogether, while age related DNA methylation changes occur they do not seem to be substantially augmented by the error prone DSB repair machinery.

## Materials and Methods

### Cell lines pre-electroporation culturing

K562 and OCI-AML3 cell lines were used in this study. The K562 Cell line was obtained from ATCC and the OCI-AML3 was a kind gift from Mark Minden’s laboratory. Both cell lines were authenticated by whole-exome sequencing and tested negative for Mycoplasma contamination. Cell lines were sub-cultured two days before electroporation in RPMI 1640 Medium containing L-Glutamine (Biological Industries, 01-100-1 A) with 10% FBS, streptomycin (20 mg/mL), and penicillin (20 unit/mL) at a density of 3*10^5^ cells/ml, at 37°C in a humidified 5% CO2 incubator.

### CRISPR/Cas9 ribonucleoprotein (RNP) cell lines editing

The sgRNA sequences for the CRISPR RNPs were chosen for the genomic loci of interest from the relevant track of the UCSC browser (https://genome.ucsc.edu/) [42]. All CRISPR/Cas9 crRNAs (S1 Table) used in the experiments were synthesized by IDT. Lyophilized sgRNAs were re-suspended in IDTE buffer (PH 7.5) to a final concentration of 100 uM. RNP complex for each reaction was generated by mixing 1.2 ul sgRNA, 1.7 ul Cas9 protein (IDT), and 2.1 ul PBS followed by incubation for 10 min at 20 degrees.

All electroporation reactions were performed using the 16-strip Lonza 4D nucleofector kit. Pre-electroporated cells were washed in PBS and spun down at 350xg for 10 min. A range of 2*10^5^ - 3*10^5^ cells per reaction were re-suspended in 20 ul SF solution (K562), SE solution (OCI-AML3) and added to the RNP complex and 1.5 ul of *ASXL1* or *SRSF2* single guide. FF-120 and EO-100 electroporation programs were used for K562 and OCI-AML3 respectively. Immediately after electroporation, pre-warmed media were added, and cells were cultured at the same conditions as the pre-electroporation culturing for four days before they were single-cell sorted.

### Generation of single cell clones

K562 cells were electroporated using sgRNA guide targeting *ASXL1* or *SRSF2* genes followed by sorting for live single-cells using BD FACSMelody™ Cell Sorter (BD Biosciences). Sorted K562 cells were plated onto 96-well plates containing 100ul/well RPMI 1640 Medium with L-Glutamine (Biological Industries, 01-100-1 A), 10% FBS, streptomycin (20 mg/mL), and penicillin (20 unit/mL). Seven days after sorting, 100 ul fresh media were added to each well. Cells were further maintained by replacing 100ul medium from each well once a week. Cell colonies were lysed 28 days after sorting for subsequent NGS sequencing. For cell lysis, cells were spun at 1000g for 5 min. Cells pellets were mixed with 50 ul of 50 mM NaOH and heated at 99 °C for 10 min. Then, the reactions were cooled down at room temperature, and 5 ul 1 M Tris PH = 8 was added to each reaction. NGS sequencing and analysis were performed as described under the following sections. Colonies containing desired repair signature for MMEJ or NHEJ indels at the genomic loci of interest were further isolated and expanded.

### Targeted sequencing of the DSB repair site

To determine the identity of the repair for each clone, we used cell lysis products that served as a template for PCR amplification and NGS library preparations. Dual indexed Illumina Libraries were generated using a two-step PCR procedure. First PCR primer prefix sequences and second PCR primer sequences were used, similar to a previously described method [29]. The targeted sequencing details are as follows: Target-specific primers were designed by Primer3plus **(http://www.bioinformatics.nl/cgi-bin/primer3plus/primer3plus.cgi)** and were ordered with the described 5’ prefixes (IDT) (Fwd – CTACACGACGCTCTTCCGATCT, Rev - CAGACGTGTGCTCTTCCGATCT). First PCR was applied to target the regions of interest. The reaction mixture was composed of a PCR-ready mix (using NEBNext® Ultra™ II Q5® Master Mix, NEB, M0544L), a cell lysis product, and a final primer concentration of 1uM, for each primer. PCR protocol was as follows: 98 °C for 30 s, followed by 40 amplification cycles of 98 °C for 10 s, 65 °C for 30 s and final elongation at 65 °C for 5 min. Following dilution of the 1st PCR products with nuclease-free water (1:1000), a 2nd PCR was performed using the Illumina sequencing primers, indexes, and adapters, under similar conditions to the 1st PCR, except for the final primer concentration of 0.5uM for each primer, and the number of amplification cycles (n=20). The primers sequences used throughout this study are provided in Table S2. The barcoded 2nd PCR products were pooled, underwent size selection (2% gel, BluePippin, Sage Science), and sequenced (paired-end, 2 × 150-bp, Miseq, illumina).

### Variant calling for the DSB repair site

Pair-end reads (2 × 150-bp) were mapped with Minimap 2.1 [43] to the hg19 genome-based targeted sequences. The resulted BAM files were further sorted and indexed using pysam 0.15.1 (https://github.com/pysam-developers/pysam). All reads from sorted BAM files were assigned to new read groups using picard 2.8.3 ‘AddOrReplaceReadGroups’ command (http://broadinstitute.github.io/picard). In order to avoid misalignments, local realignment was performed using GATK 3.7 ‘RealignerTargetCreator’ and ‘IndelRealigner’ [44]. Mpileup files were generated by samtools 1.8 followed by SNVs and small indels detection using varscan 2.3.9 ‘pileup2cns’ command to generate VCF files containing consensus variant calls [45]. Frequencies were calculated as the sum of modified reads divided by the mean depth per experiment.

### Methylation quantification with Targeted bisulfite sequencing

Genomic DNA was extracted from the desired expanded single cell clones using Qiagen DNeasy columns and measured with nanodrop. An amount of 200 ng of genomic DNA from each sample underwent bisulfite conversion with Zymo DNA Methylation Gold Kit (Zymo Research). The converted DNA served as a template for PCR amplification and library preparations as in the former section. Target-specific primers were designed manually with Tm adjustment using the Thermo Fisher Tm calculator (https://www.thermofisher.com/) and were ordered with the described 5’ prefixes (IDT).

First PCR was applied to target the regions of interest. The reaction mixture was composed of a GoTaq® Reaction Buffer (Promega, M792A), PCR Nucleotide Mix, GoTaq® G2 DNA Polymerase (Promega, M784A), a bisulfite converted DNA product, and a final concentration of 1uM for each primer. PCR protocol was as follows: 95 °C for 3 min, followed by 36 amplification cycles of 95 °C for 20 s, 56 °C for 30 s and 72 °C for 90 s, the final elongation was at 72 °C for 5 min. The PCR product was processed as described for the DSB repair targeted sequencing.

Methylation was quantified based on the alignment of bisulfite converted products to hg19 based genomic sequences where all the cytosine bases were replaced with thymidine except in CpG sites. Sites with less than ten reads of coverage were discarded. The methylation rate was calculated for each CpG site in the relevant section of the Hg19 genome, as the compliment to the ratio between converted base (‘T’) to the sum of converted and non-converted bases (‘C’+’T’), 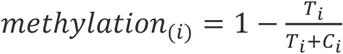.

---

**Supplementary Table 1.**
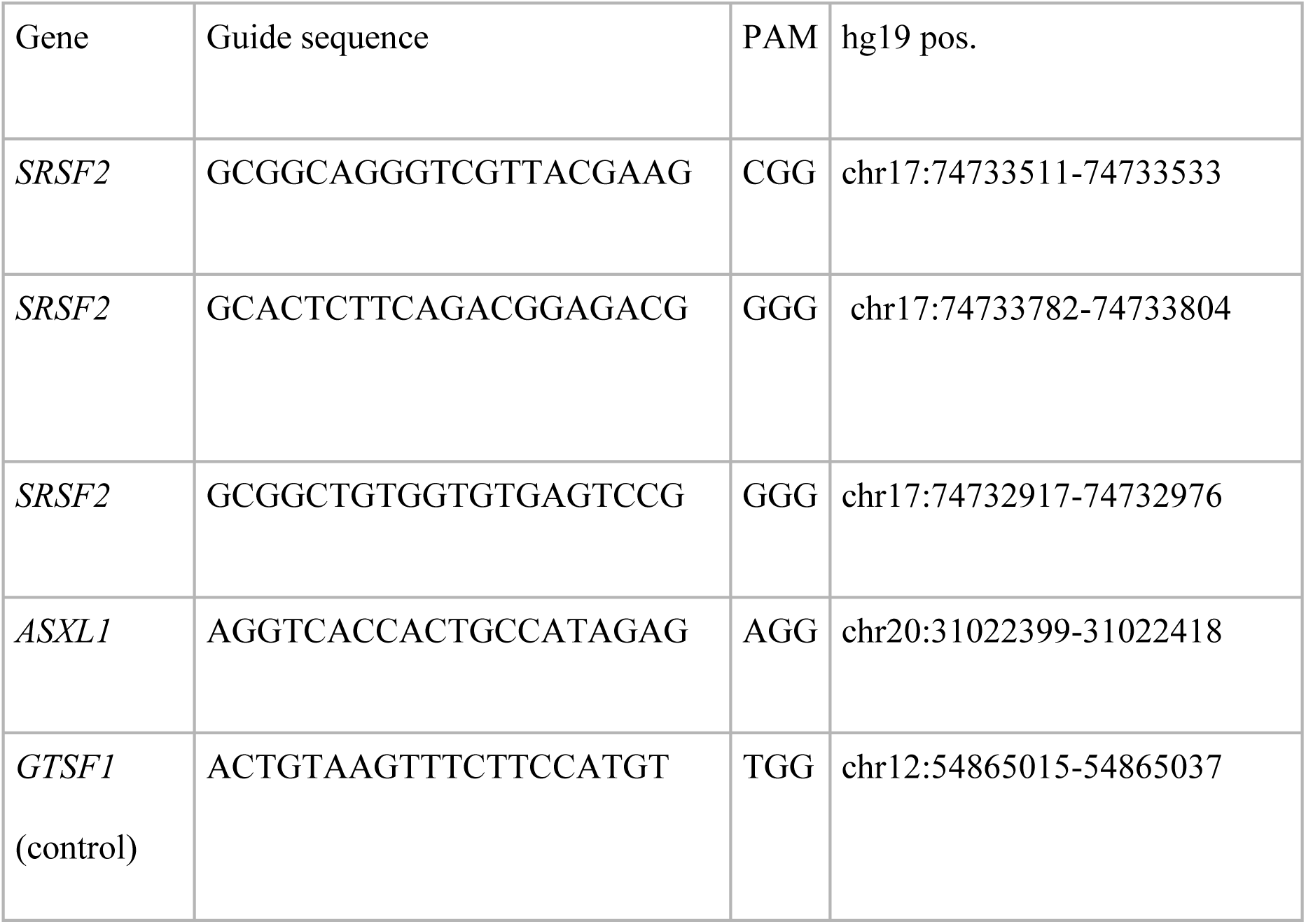
*ASXL1* and *SRSF2* and control gRNA sequences.

**Supplementary Table 2.**
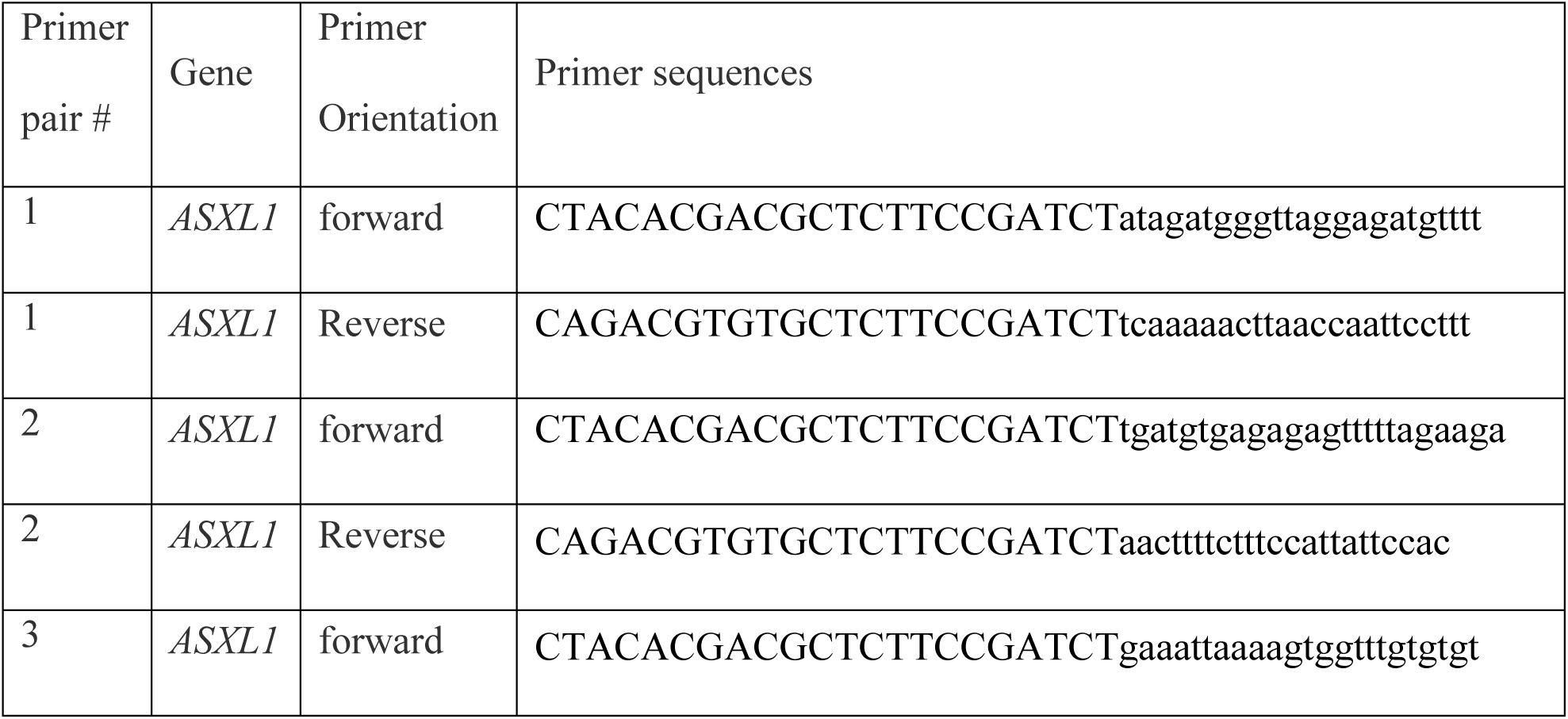
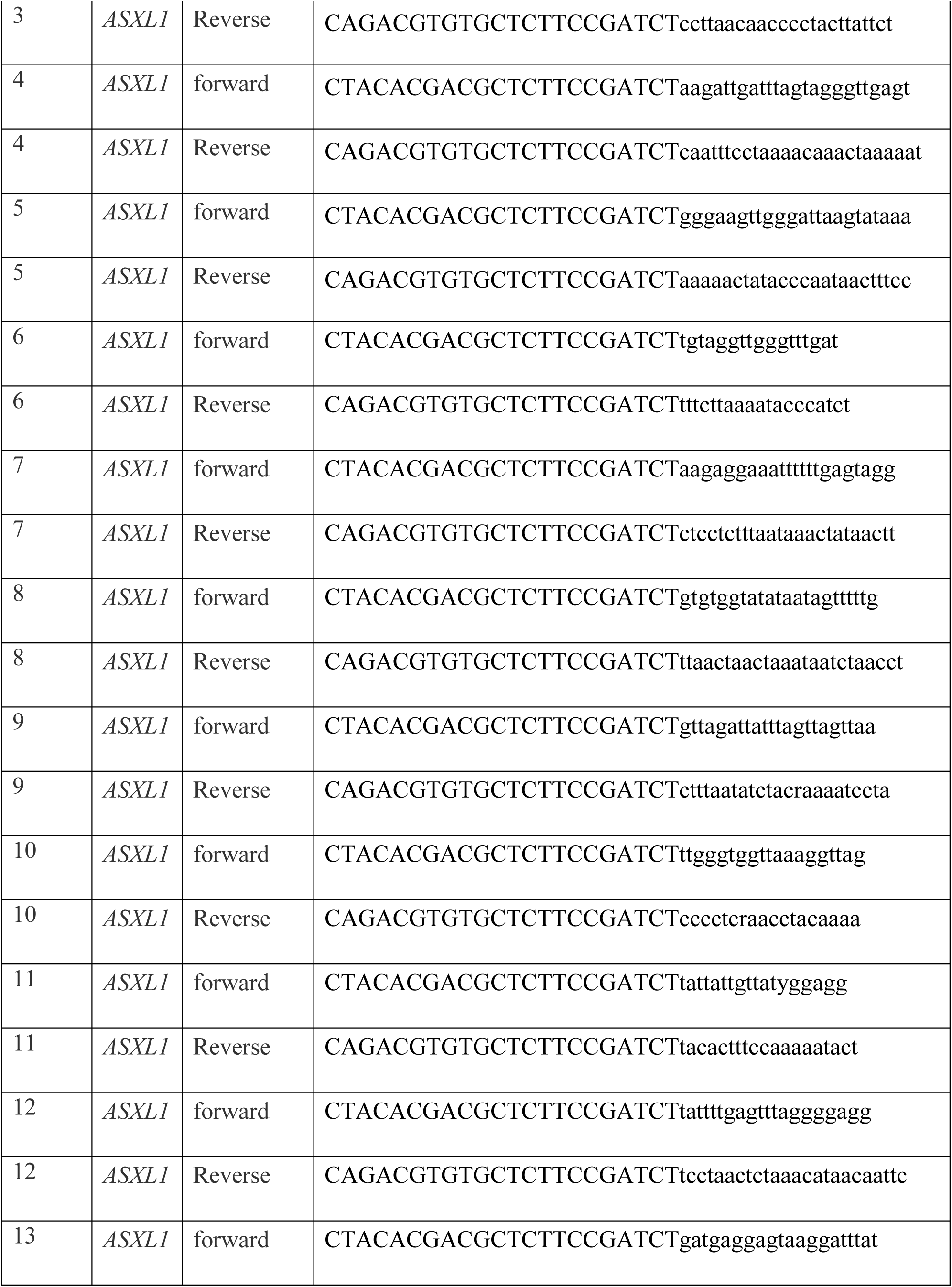
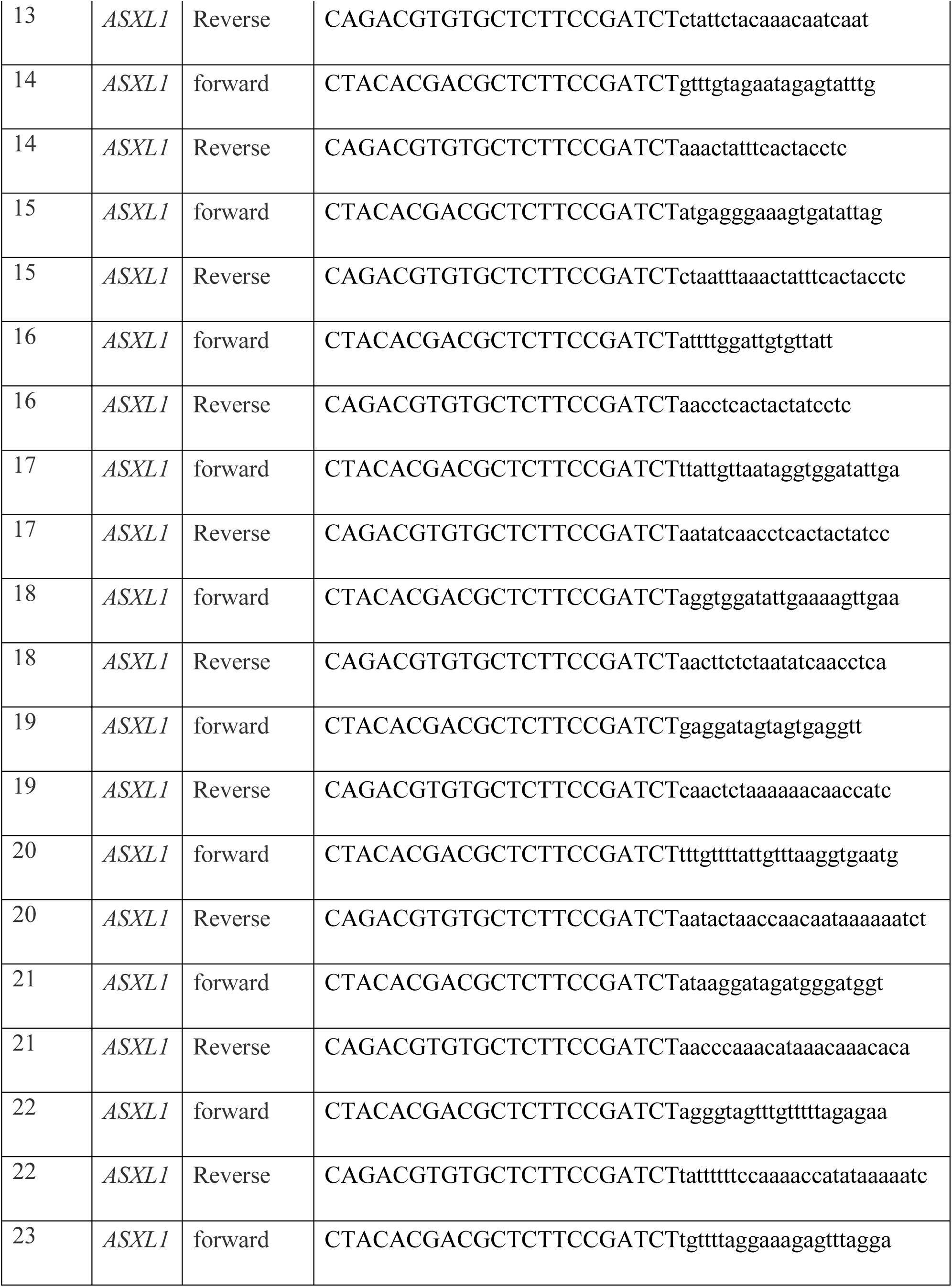
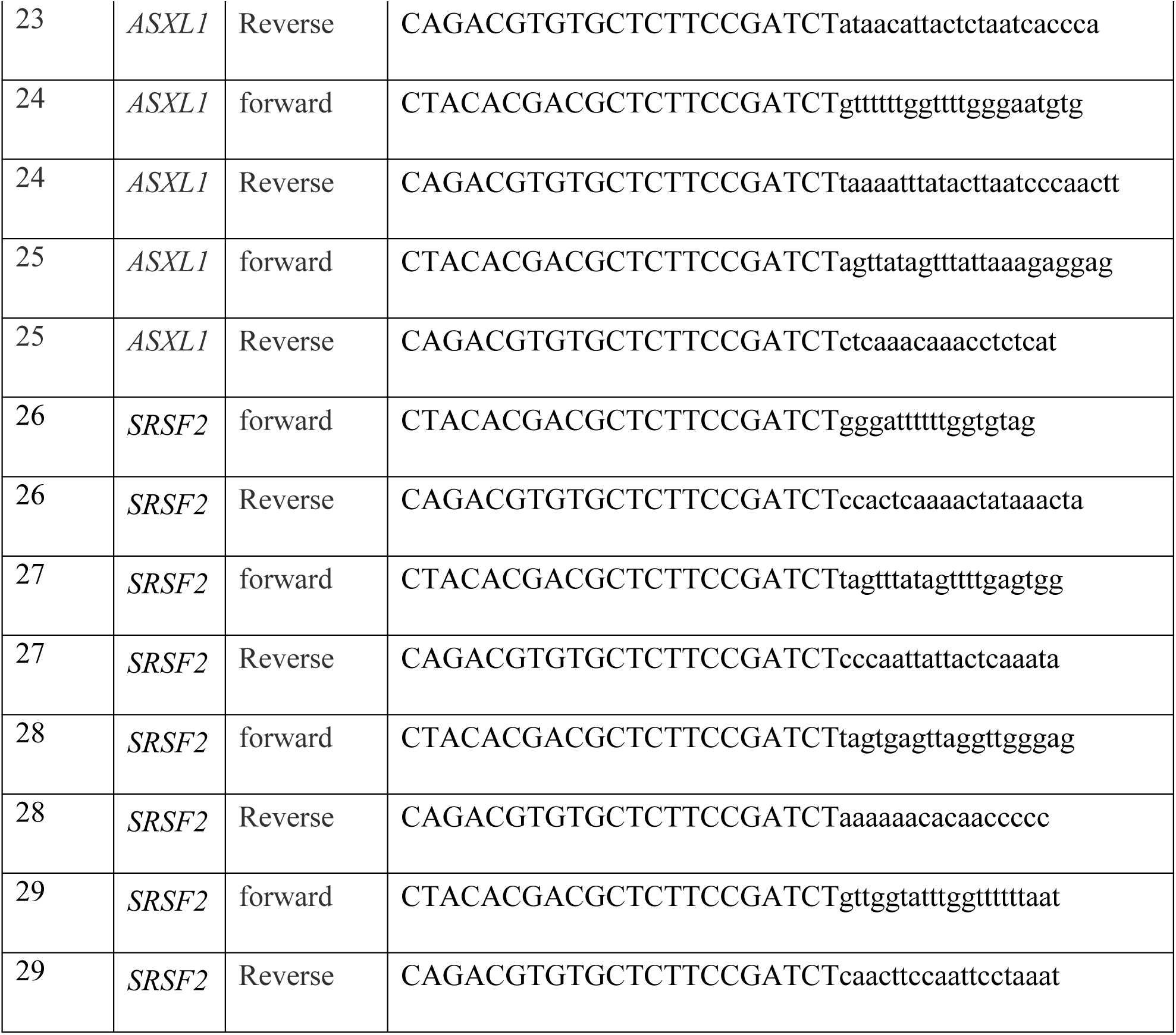
A list of primers that were used for repair pathway determination and methylation quantification. Capital letters indicates the sequences used in the second PCR.

**Supplementary Table 3 (CSV file)**. List of genetic changes found in the single cell clones.

**Supplementary Table 4 (CSV file)**. Methylation data found in K562 single cell clones in CpG sites around the DSB site of *ASXL1*.

**Supplementary Table 5 (CSV file)**. Methylation data found in OCI-AML3 single cell clones in CpG sites around the DSB site of *ASXL1*.

**Supplementary Table 6 (CSV file)**. Methylation data found in K562 single cell clones in CpG sites around the DSB site of *SRSF2*.

**S1 Fig.**
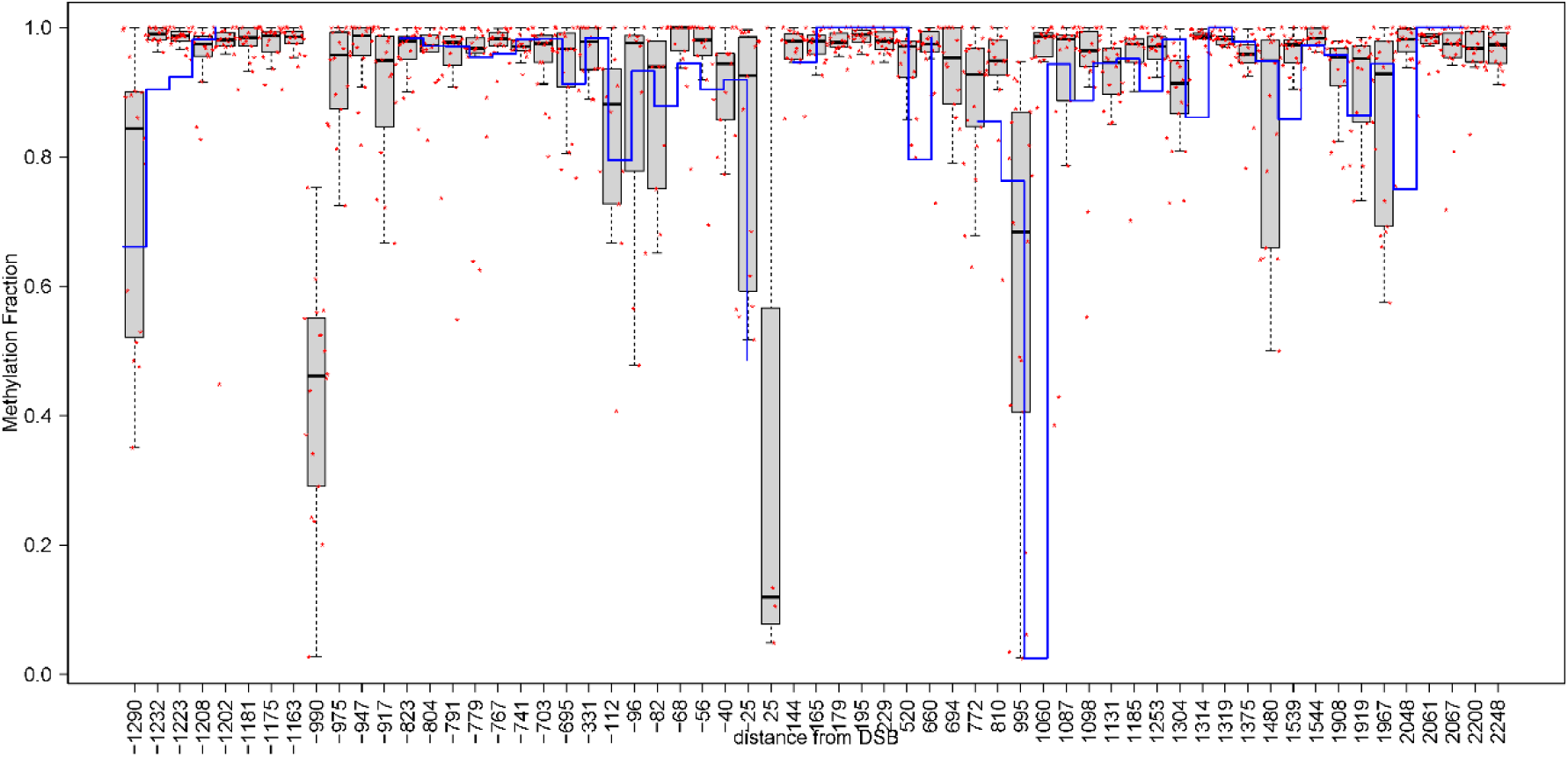
Methylation variability in non-edited control single cell clones. Boxplots represent the distribution of methylation level as measured in control single cell clones (red dots) around *ASXL1* (X-axis coordinates relative to the DSB site in the non-control cells). The level of methylation as was measured in bulk (blue line) is high and CpG sites with lower methylation level in the bulk represent variability rather than low level of methylation.

**S2 Fig.**
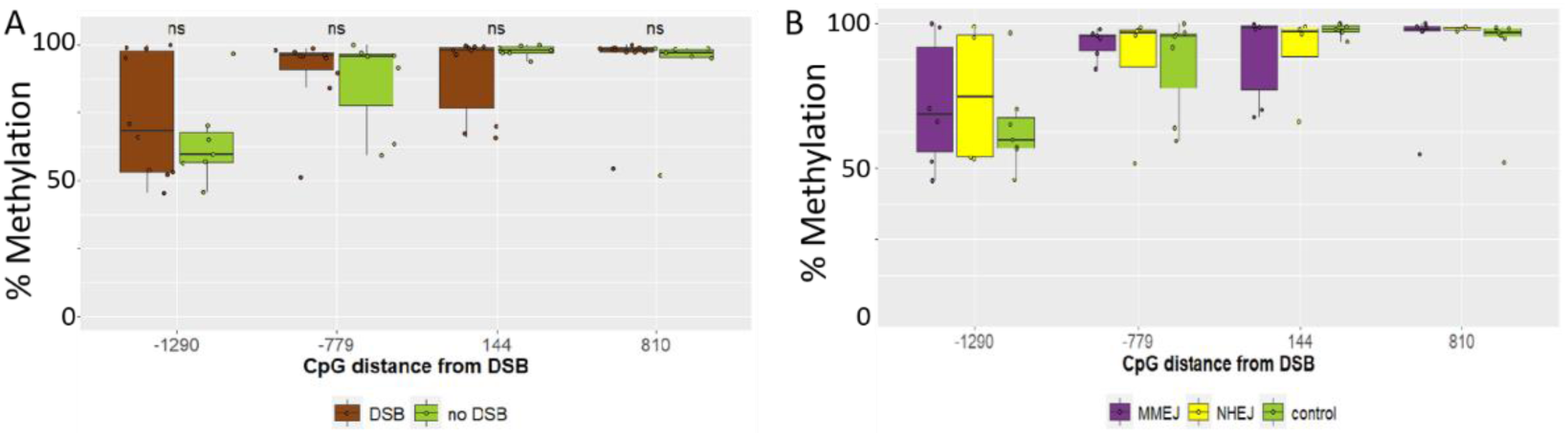
Selected sites to demonstrate methylation stability in OCI-AML3 single cell clones following DSB in ASXL1. Comparison of methylation levels in selected highly variable CpG sites for (A) all DSB induced outcomes (brown) to controls (green) or (C) according to the mechanism of repair: MMEJ (purple), NHEJ (yellow), and controls (green). No Significant difference was detected.

**S3 Fig.**
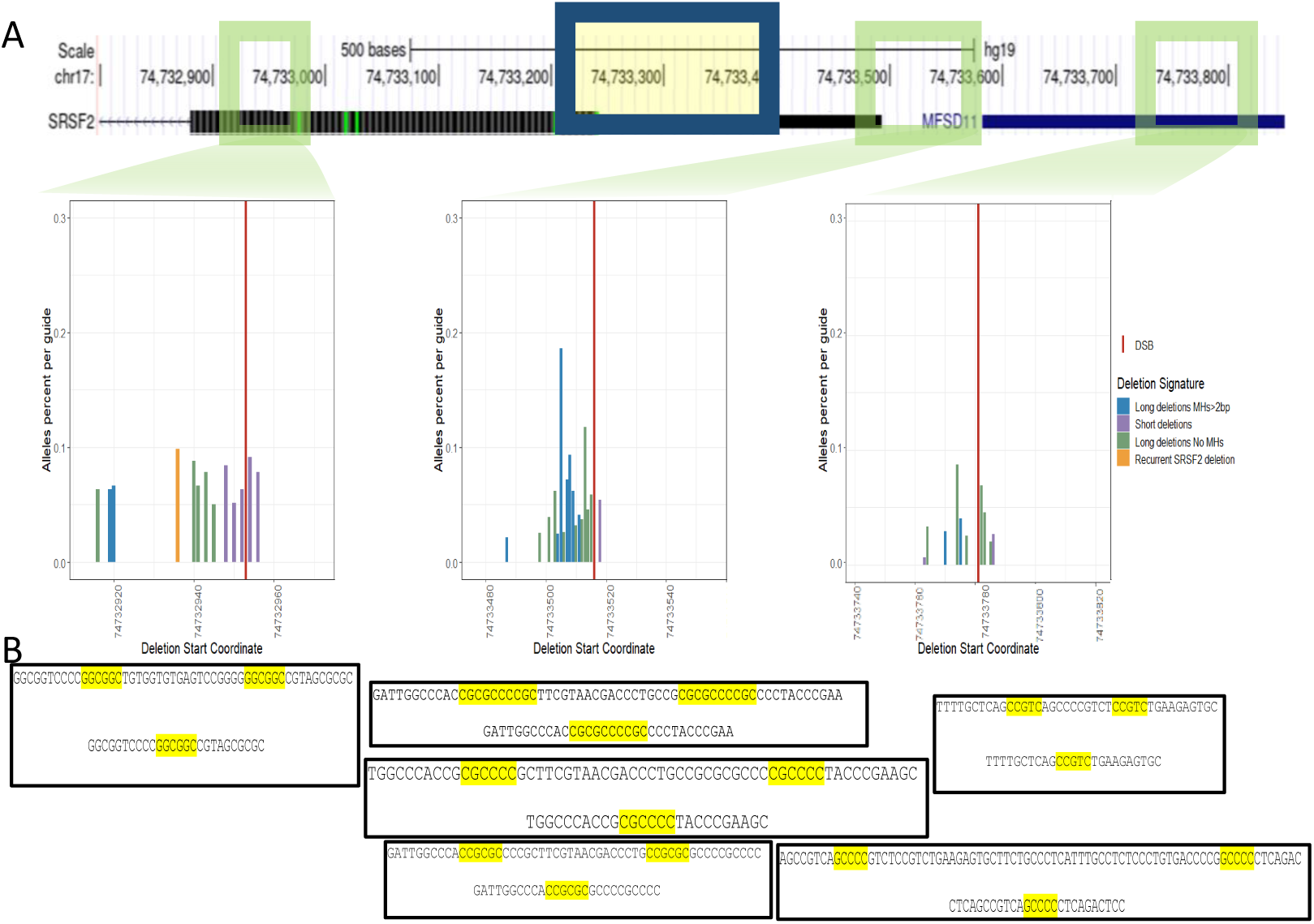
Characterization of the DSB repair in SRSF2 gene body and promoter of *SRSF2*. **(A)** Representation of the *SRSF2* region with DSB locations (green frames). The repair outcome of the three gRNAs is below each frame. Dark blue frame indicate the region where the methylation was quantified. **(B)** Selected clones to represent the MMEJ repair pathway with homolog sequence (yellow shading) greater than 5 bp. The sequences appear below the gRNA used for the DSB induction.

